# Protein design under competition for amino acids availability

**DOI:** 10.1101/331736

**Authors:** F. Nerattini, L. Tubiana, C. Cardelli, V. Bianco, C. Dellago, I. Coluzza

## Abstract

Understanding the origin of the 20 letter alphabet of proteins is a long-lasting biophysical problem. In particular, studies focused extensively on the effect of a reduced alphabet size on the folding properties. However, the natural alphabet is a compromise between versatility and optimisation of the available resources.

Here, for the first time, we include the additional impact of the relative availability of the amino acids. We present a protein design scheme that involves the competition for resources between a protein and a potential interaction partner that, additionally, gives us the chance to investigate the effect of the reduced alphabet on protein-protein interactions. We identify the optimal reduced set of letters for the design of the protein, and we observe that even alphabets reduced down to 4 letters allow for single protein folding. However, it is only with 6 letters that we achieve optimal folding, thus recovering experimental observations.

Additionally, we notice that the binding between the protein and a potential interaction partner could not be avoided with the investigated reduced alphabets. Therefore, we suggest that aggregation could have been a driving force for the evolution of the large protein alphabet.

## INTRODUCTION

The amino acid alphabet encoding the protein function is common to all living organisms and is the result of millions of years of evolution. It is composed of 20 letters, in contrast to the ones of other biopolymers, such as DNA and RNA, which possess 4 letters only. Such a large alphabet gives to proteins the vast variety of configurations and functions that we know so far.

The advent of artificial protein evolution (also known as protein design) (1–16) opens the possibility to address fundamental questions about the nature of the amino acid alphabet (17–20). One of the questions that mostly attracted the attention of the scientific community was about the universality of the 20 letters. Why 20 and not less? Could it be possible to design proteins to fold using a reduced alphabet? The early work on protein design with alphabets of different sizes was carried out for protein lattice models in which the protein chain is constrained to be on a cubic lattice. With such models it was possible to design heteropolymers with a large variety of alphabets defined by the amino acid interactions (21–30). It became rapidly apparent that even in such simplified systems it is necessary to have a minimum number of residue types to encode the target configurations (31). Moreover, such simple models allowed to explore the related question on how the alphabet size influences protein-protein interactions (32–35). Finally, works done on realistic models, offer substantial evidence that protein design with a minimalistic alphabet is possible (36–40). In particular, statistical analysis of protein databases demonstrated that a considerable fraction of the information encoded in natural proteins could be packed into smaller efficient alphabets of just 5 residue types (36, 38, 41–44). However, all the mentioned studies completely neglected the possibility that a competition for the availability of amino acids may have played a role in the evolution of the protein alphabet size.

In this work, we devised a design strategy which not only include such a competition, but also check the effect of a reduced alphabet on protein folding and protein-protein interaction. We consider systems composed of the natural protein G (PDB ID: 1PGB, already successfully redesigned with several protein models (3, 7)) and a potential binding partner (a mould of a part of protein G, that mimics with a surface-like shape a potential binding site of a larger protein). Both protein and binding partner are represented with the Caterpillar coarse-grain model, which has been successfully tested to design and refold natural and artificial proteins (7, 9). We simultaneously design the sequences of the protein-partner system according to different optimisation pathways, namely optimal folding for the protein G and just optimal interaction with the solvent for the binding partner.

The adopted design scheme leads to a competition for the available amino acids, since the protein and the surface cannot use all the 20 amino acids simultaneously for the sequences optimisation. In practice, although the whole alphabet accessible by the binary system is still composed of 20 amino acids, the condition that we impose to the design procedure are such that the protein will have a smaller alphabet available to optimise the folding: the larger the binding surface, the stronger is its effect on the segregation.

Such a procedure allow us to control the strength of the competition as a function of the size and geometry of the binding partner. We obtain the optimal protein alphabet with the minimum number of letters, without the need of imposing the composition of the protein. Additionally, by having a protein surface system, we can explore the effect of the alphabet segregation on the aggregation in different protein-surface binding scenarios. The results show that for the folding of a small protein the minimum number of amino acid types needed is just 4. However, such a small alphabet compromises the heterogeneity of the protein-protein interactions (21, 29, 33–35) and binding cannot be avoided.

This result has interesting implications towards the understanding of the evolution of protein sequences and structure when the amino acid availability is taken into account. In fact, living systems are under constant pressure for using the smallest variety of amino acids as possible, e. g. to limit the resources needed to construct specialised tRNA molecules necessary for the translation process (45). Hence, it is reasonable to assume that during the early stages of life, the protein capable of being designed with a smaller alphabet could have been advantageous. If protein aggregation was not crucial at that stage, then our results demonstrate that protein-based life could have started with alphabets size compatible with the one of DNA and RNA. On the other hand, the simple condition of avoiding protein aggregation could be a strong driving force against alphabet segregation.

The structure of the article is the following: firstly we discuss the modelling procedure to construct a test system for protein-protein interactions that allow us to perform a protein design under competing conditions for amino acid availability. We then describe the computational method for studying design and folding of such a test system. In the central part of the article, we show the results regarding the reduced alphabets, the folding properties of the protein alone and in the presence of a binding partner. In the last part, we highlight the main conclusions of our investigation.

## MATERIALS AND METHODS

### Protein model

The Caterpillar protein model reduces the amino acid structure and represent it by the backbone atoms: *C*, *O*, *C*_*α*_, *N*, *H* (Fig. 1(a)). The intramolecular energy for a protein of length *N* has the form:

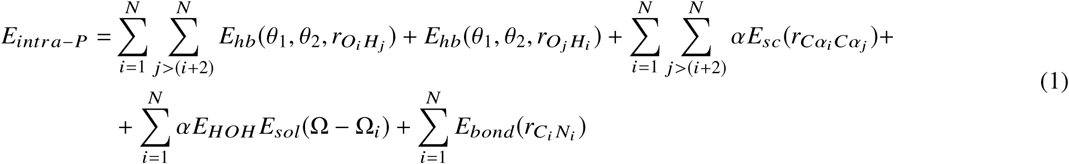

where *α* and *E*_*HOH*_ are added to balance the relative weight of different energy terms. *E*_*hb*_(*θ*_1_, *θ*_2_, *r*_*OH*_) is a 10 – 12 Lennard-Jones potential commonly used to represent hydrogen bonds (46):

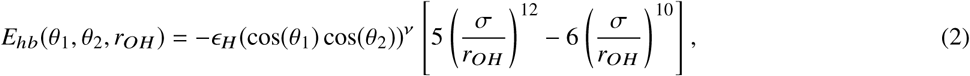

being *r*_*OH*_ the distance between the hydrogen atom of the amide group and the oxygen atom of the carboxyl group of the main chain; *θ*_1_, *θ*_2_ the angles between the atoms *COH* and *OHN* respectively (Fig. 1(a)), and account for the hydrogen bonds directionality; ν = 2; σ = 2 Å and ϵ_*H*_ = 3.1*K*_*B*_*T*.

**Figure 1:**
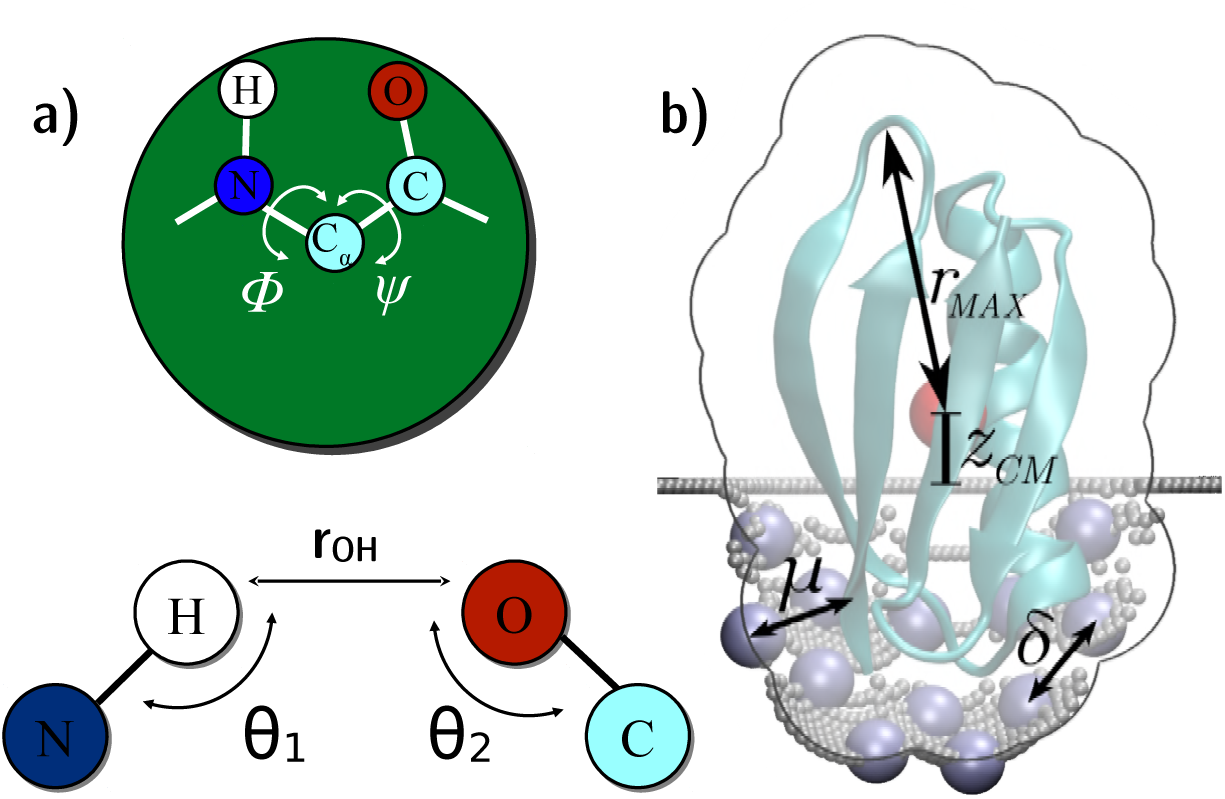
**a)** Caterpillar protein model: the green circle represents the amino acid self-avoidance volume, which has a radius of 2 Å and is centred on the position of the *C*_*α*_ atom. Each amino acid is represented through backbone atoms only. Side chain-side chain interactions are represented via a square-well-like potential (Eq. 3).For the hydrogen bonds, the model include a 10 12 Lennard-Jones potential, that is a function of the distance *r*_*OH*_, and tunes it with a multiplicative factor that involves the angles *θ*_1_ and *θ*_2_, so to account for the directionality of the bond. **b)** Artificial binding site parameters. Gray dots are self-avoiding beads; blue spheres are *C*_*α*_ atoms of the binding site. The red spot represents the centre of mass (CM) of the protein, *z*_*C M*_ is its height with respect to the *z* = 0 plane and *r*_*M AX*_ is the maximum CM-*C*_*α*_ distance. *µ* is the minimum *C*_*α*_ protein-*C*_*α*_ surface distance, i.e. the value one has to use to inflate the protein for the binding site shaping. *δ* is the minimum distance between two activated *C*_*α*_ of the binding site.

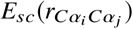 mimics the side chain-side chain interaction via a smoothed square-well-like isotropic potential:

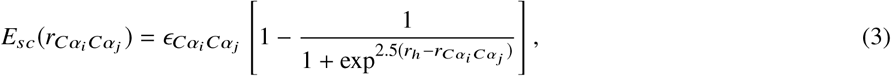

where 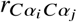 is the distance between the *C*α atoms and *r*_*h*_ = 12 Å. Eq. 3 is a sigmoid function with a flex at 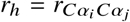, where 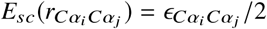. The terms 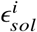 are the residue-residue interaction parameters of the interaction matrix and their value is taken from Tab. S1 of Ref. (9).

*E*_*sol*_(Ω - Ω_*i*_) is an implicit solvent energy term that acts as an energy penalty if a hydrophobic (hydrophilic) amino acid is exposed (buried), and has the form:

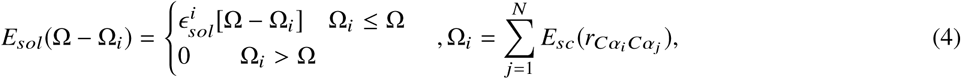

where Ω = 21 is a threshold for the number of contacts in the native structure above which the amino acid is considered to be fully buried, and 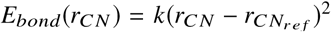 is the Dolittle hydrophobicity index (47).

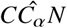 is a harmonic bonding term with elastic constant *k* = 20 *K*_*B*_*T* Å^−2^, that keeps fixed the distance *r*_*C N*_, along with the *Cĉ*_*α*_*N* backbone angle.

For more details about the model see Ref. (7, 9). For the binding site amino acids, instead, we use a single atom representation and make use of the *E*_*sol*_ and *E*_*sc*_ terms only.

### Binding site model

The binding site is modelled as a fixed layer of amino acids, idealizing a small pocket on the surface of a much larger globular protein. To represent the artificial binding site, we generate a mould by pushing the protein on a mesh of self-avoiding beads initially lying flat on the *z* = 0 plane, as illustrated in Fig. 1(b). The high density of beads in the mesh prevents the protein from passing through it. A finite number of *C*_*α*_s, the type of which is assigned during the design procedure, are distributed on the surface binding site, to model its protein nature. We refer to the latter atoms as *C*_Surf_.

There are three important parameters in the modelling procedure: *ζ*, the reduced centre of mass (CM) height with respect to the *z* = 0 plane; *µ*, the minimum *C*_*α*_ protein-*C*_*α*_ surface distance; and *δ*, the gap between two *C*_*α*_s of the binding site. To define the reduced height *ζ*, we first identify *r*_*M AX*_, i.e. the maximum protein radius with respect to the CM (*r*_*M AX*_ 16 Å for protein G), and then use it to normalise the height *z*_*C M*_ of the CM from the *z* = 0 plane: 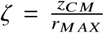. Hence, we push the protein into the *z* = 0 plane until we reach the desired *ζ* value. The mesh is pushed downwards accordingly, keeping every point at the minimum distance between all atoms of the protein and the binding site at *µ* = 13 Å (see Fig. 1(b)). The value *µ* = 13 Å ensures a low influence of the binding site on the protein design regarding the protein-binding site energy *E*_*sc*_ (Eq. 3). We construct systems with different *ζ* values, and for each one of them, we perform a rotational analysis by keeping the position of the CM fixed and by shaping the mesh for various orientations of the protein until we reach the maximum surface area of the binding site. It is important to notice that the surface area decreases with the increase of *ζ*.

Having obtained the geometrical shape of the pocket, we proceed to “activate” a subset of its beads by assigning to them a *C*_*α*_ nature. These beads constitute the *C*_*sur f*_ set. The activated beads are homogeneously distributed in the pocket and are always separated by a distance of *δ* = 5 Å from each other. The value of *δ* is derived from the typical nearest distance between two residues in natural proteins, as explained in Fig. S8 of the Supplementary Information (SI).

Since the amino acids in the *C*_Surf_ set are represented by *C*_*α*_ atoms, the protein-binding site interaction energy includes the Caterpillar *E*_*sc*_ and *E*_*sol*_ terms only (see Eqs. 3 and 4). Given that our binding site representation is limited to the surface residues, such an approximation affects the correct evaluation of the binding site solvent exposure term *E*_*sol*_. Since in turns *E*_*sol*_ will influence the protein binding properties, we add an offset to the number of contacts for each amino acid of the binding site (∼ 6), to correctly account for the solvent exposure term *E*_*sol*_ of the binding site amino acids. The offset was calculated in such a way that the maximal amino acid exposure is compatible to the one of natural proteins (e.g. the protein G itself described in Fig. S10 of the SI). Finally, since the binding site conformation is fixed, we neglect all the interactions between all the pairs of the *C*_*sur f*_ set.

For more information about the algorithms used in the binding site modelling procedure, see the section *Binding site modelling* of the SI.

### Design

To investigate the sequence space, we perform Virtual Move Parallel Tempering (48) (VMPT) Monte Carlo simulations with swap and single point amino acid mutation moves along the sequence, a procedure that has been already successfully employed in protein design (7, 9, 16). We simulate the same system at different temperatures, and swap sequences between the replicas on the fly, thus enhancing the overcoming of energy barriers and, therefore, improving the sampling. Moreover, at each temperature, we collect statistics using the information coming from all other replicas, according to the virtual move scheme described in Ref. (48). In our implementation we simulate 16 replicas with a set of temperatures (10.000; 5.000; 2.000; 1.000; 0.500; 0.333; 0.250; 0.200; 0.167; 0.143; 0.125; 0.111; 0.100; 0.091; 0.083; 0.077) in units of *K*_*B*_. We also consider the protein-binding site as a single object, frozen in the target conformation generated via the binding site modelling described above. The sequences of the protein and the binding site are optimised jointly. The best candidate sequences for the folding are the ones which minimise the energy of the target structure and maximise the number of permutations *N*_*p*_, given by:

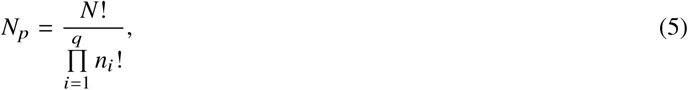

to increase the composition heterogeneity along the joint protein-binding site sequence. In Eq. (5) *q* = 18 is the alphabet size (proline and cysteine are not included in the design, due to their peculiar role in protein structure, which is beyond the scope of our model); *N* is the total number of monomers in the protein-binding site system; *n*_*i*_ is the number of monomers of type *i* in the joint sequence.

We enhance the sampling by introducing an adaptive bias potential *W* [*E*, ln *N*_*p*_, *T*] over two collective variables *E* and *N*_*p*_. *W*[*E*, ln *N*_*p*_, *T*] is used to bias the acceptance probability of each Monte Carlo mutation move. Therefore, each Monte Carlo step is accepted with a probability

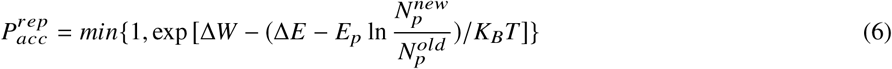

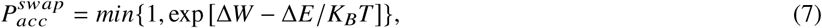

associated to the amino acid replacement and amino acids swap moves, respectively. In the latter equations Δ*W*, Δ*E* and ln 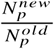 refer to the differences of the bias potential, the energy and the number of permutation between the new configuration and the old one; *E*_*p*_ = 20 *K*_*B*_*T* is a scaling factor set to a value high enough to generate highly heterogeneous sequences.

### Folding

We adapted the Monte Carlo folding procedure for a single protein, described in Ref. (7, 9), to handle a system composed of interacting partners. The simulation is performed in a cubic box (as shown in Fig. S11) containing two replicas of the binding site (box side ∼ 360 Å), that are the mirror image of the other with respect to the *xy* plane passing through the centre of the box.

The binding sites are frozen in the box, and the protein starts the folding from a fully stretched conformation. An impenetrable slab region is defined between the binding sites to prevent the protein to approach them from the convex side instead of the concave one generated with the moulding procedure.

We sample the protein conformations using crankshaft, pivot, rotation, translation and mirroring moves, to let the protein reorient, diffuse in the box and switch from right to left handed conformations. The simulation is performed in parallel at 32 different temperatures and the replicas exchange information through the VMPT bias scheme (48). The set of reduced temperatures (8.5; 7.8; 7.2; 6.6; 6.0; 5.4; 4.9; 4.6; 4.3; 3.9; 3.5; 3.1; 2.8; 2.55; 2.35; 2.2; 2.05; 1.9; 1.75; 1.6; 1.45; 1.3; 1.1; 1.0; 0.95; 0.92; 0.88; 0.85; 0.82; 0.79; 0.76) is chosen so to observe the protein repeatedly detaching from the binding site.

Each replica sampling is enhanced by a bias potential depending on two collective variables, namely the distance root mean square displacement within the protein *DRMSD*_*intr a*_, and between protein and binding site, *DRMSD*_*inter*_. The *DRMSD* is a measure of the distance of a configuration from a target structure:

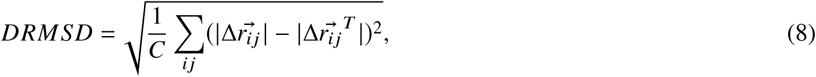

where the sum runs over the *i j* pairs in contact in the target structure (namely the native contacts, identified using a cut off of 17 Å according to previous studies (7, 9)), 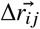 is the distance between the residues *i* and *j* belonging to the same (*DRMSD*_*intr a*_) or different (*DRMSD*_*inter*_) proteins, and 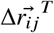 is the same distance calculated over the target structures. Accordingly, the *DRMSD*_*intr a*_ is computed adopting the native and isolated protein conformation as a target structure, while the *DRMSD*_*inter*_ is computed using as a target structure the native protein bound to the binding site. The normalisation factor *C* is the number of native contacts for *DRMSD*_*inter*_ and twice the number of native contacts for *DRMSD*_*intr a*_. 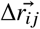 for *DRMSD*_*inter*_ is evaluated with respect to the binding site closest to the protein for each *i j* pair.

## RESULTS AND DISCUSSION

The purpose of the present work is to design proteins with the optimum reduced amino acid alphabet, to check both their folding abilities and their aggregation behaviour in the presence of a possible binding site. To this aim, we chose a value of *µ* = 13 Å that minimises the optimisation of the protein-binding site interaction (see section *Binding site model*), thus leading to the design problem described in the following.

The globular structure of the protein leads to a complex design optimisation problem since every residue is in contact with many others and the water exposure profile varies from the surface to the buried residues. On the other hand, the residues of the binding site possess few contacts; these weakly interacting residues are mainly optimised for the exposure to the solvent. Since the coupling between these two optimisation procedures is only through Eq. (5), i.e. the condition of maximizing the composition heterogeneity, the design leads to the spontaneous segregation of a reduced alphabet on the protein sequence, keeping the variability high on the one of the binding site.

We investigate four systems differing in terms of *ζ* = (0.20, 0.40, 0.60, 0.80), thus leading to a surface area = (4717.5, 3842.2, 3051.5, 2320.5) Å^2^ and *C*_Surf_ = (158, 127, 100, 78) residues respectively. For all scenarios, we generated a large number of sequences that we group in solution basins. In Fig. 2 we plot the distribution of the Hamming distances calculated for all the possible pairs between the basins. The latter is measured using the Hamming distance that determines the difference between two sequences of equal length by counting the number of substitutions needed to make them coincide. It is often more convenient to normalise the Hamming distance by dividing by the protein length *N*. For all scenario we observe large number of solutions that differ for more than 50% of the residues. However, *ζ*= 0.60 and 0.80 have also a significant number of related solutions with sequences differing for 20% only (see Fig. 2(f)). To check that nevertheless the *ζ*= 0.60 and *ζ*= 0.80 basins are distinct, we also calculated the self-overlap distributions for *ζ*= 0.60 and *ζ*= 0.80 (see Fig. S9 in the SI) and we obtained profiles that are different from each other. More importantly, they differ from the one plotted in Fig. 2(f), demonstrating that although there are similarities, the two basins are distinct.

**Figure 2:**
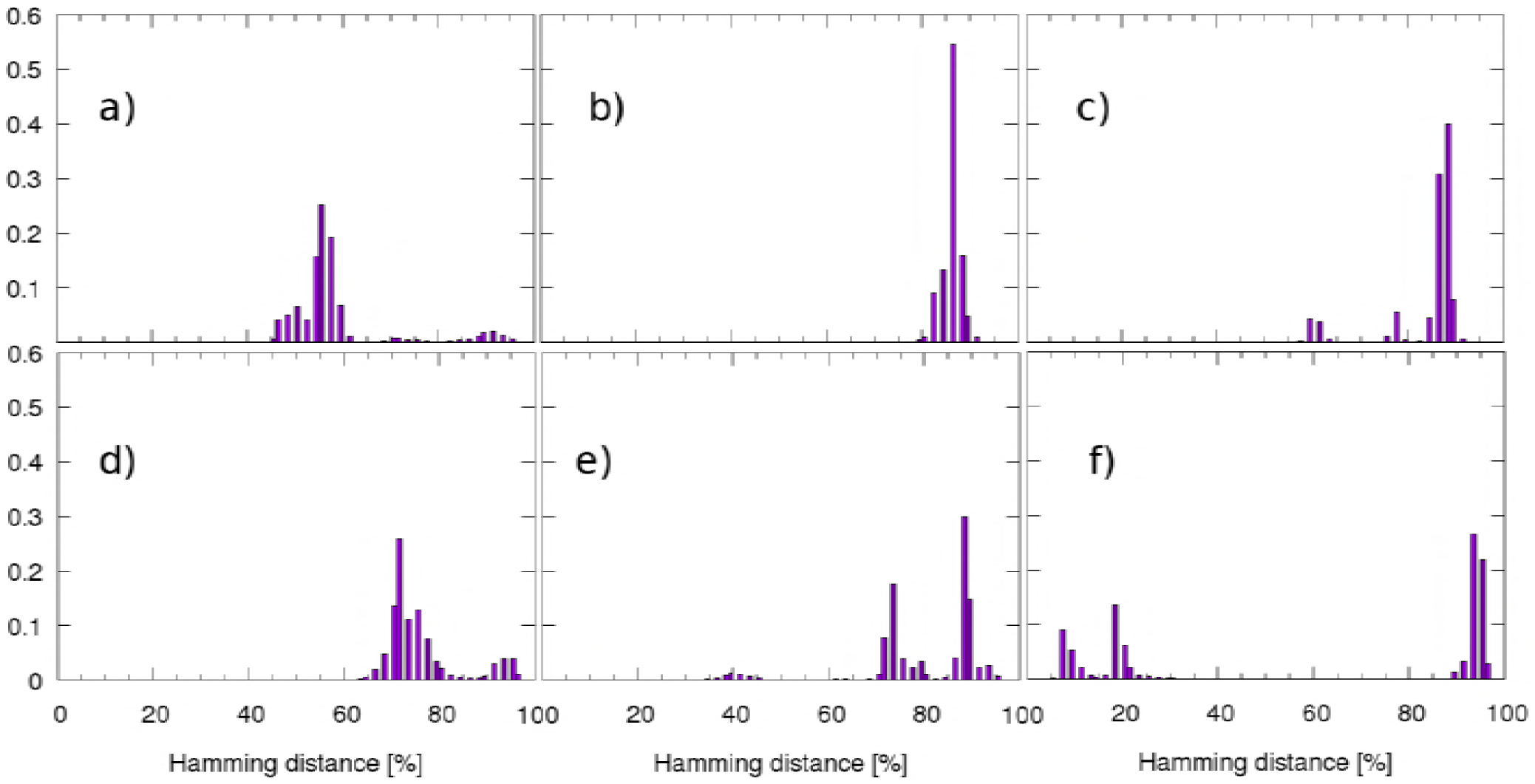
Histograms obtained by evaluating the Hamming distance (% relative to the chain length) between all possible pairs of sequences chosen selecting 200000 solutions around the design free energy minimum of systems *A* and *B*, corresponding to the *ζ* values specified in the following. a) *A*: *ζ*= 0.20; *B*: *ζ*= 0.40 b) *A*: *ζ*= 0.20; *B*: *ζ*= 0.60 c) *A*: *ζ*= 0.20; *B*: *ζ*= 0.80 d) *A*: *ζ*= 0.40; *B*: *ζ*= 0.60 e) *A*: *ζ*= 0.40; *B*: *ζ*=0.80 f) *A*: *ζ*= 0.60; *B*: *ζ*= 0.80

A crucial feature of the distributions in Fig. 2, are the peaks at high Hamming distances. They demonstrate that the protein-binding site coupling induces the design process to explore distinct solution basins. Given that protein and binding site are energetically very weakly coupled (evident from the low value of *E*_*sc*_-inter that ranges from –1.4 to –2.7 *K*_*B*_*T*), the influence of the binding site can be attributed to the maximisation of the letter permutations expressed by Eq. (5). Such coupling enforces an increasing competition between binding site and protein for the alphabet available while increasing the binding site size.

In Fig.3 we plot the occurrence frequency of each amino acid type in four sequences selected with highest permutation number and lowest energy among the ones in the solution basins (shown in Tab. 2 and Tab. 1).

**Table 1:** Hamming distance [%] between designed protein sequences.

| $\zeta$ | 0.20 | 0.40 | 0.60 | 0.80 |
| --- | --- | --- | --- | --- |
| 0.20 | 0 | 61 | 87 | 86 |
| 0.40 |  | 0 | 84 | 86 |
| 0.60 |  |  | 0 | 27 |
| 0.80 |  |  |  | 0 |

**Table 2:**
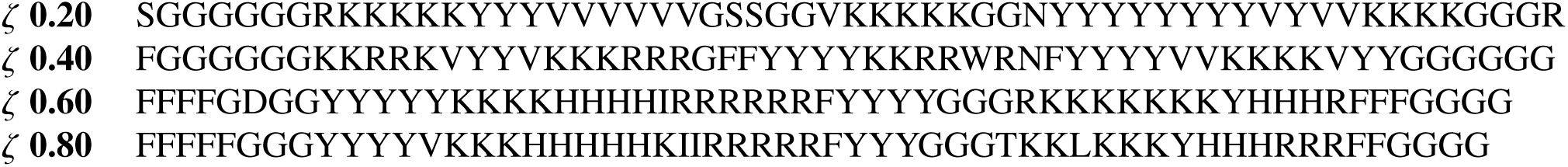
Protein G designed sequences for the investigated systems.

Given that the Caterpillar alphabet is composed by 18 letters, the frequency of each amino acid in a maximally heterogeneous sequence would be 1 18 ∼ 6%. Therefore, we set a threshold at a slightly higher value=10%, to safely discern the contribution of dominating amino acids from the random occurrences. From the plot it becomes apparent that the protein designed sequence has a restricted composition of amino acids leading to a segregated effective protein alphabet. We observe that the effective alphabet grows from 4 to 6 letters going from larger (*ζ*=0.20 and 0.40) to reduced binding sites (*ζ*=0.60 and 0.80) respectively. It is interesting to notice that the alphabets are made of amino acids with an average attractive pair-interaction energy and high variability in terms of the residue-solvent interactions (see Table S1 in Ref (9)). Moreover, the alphabets differ from each other, and for each scenario, the protein amino acids are not present in the corresponding binding sites sequence (see Fig. 3). The latter finding shows that the design process indeed mimics a process under competition for available amino acids.

**Figure 3:**
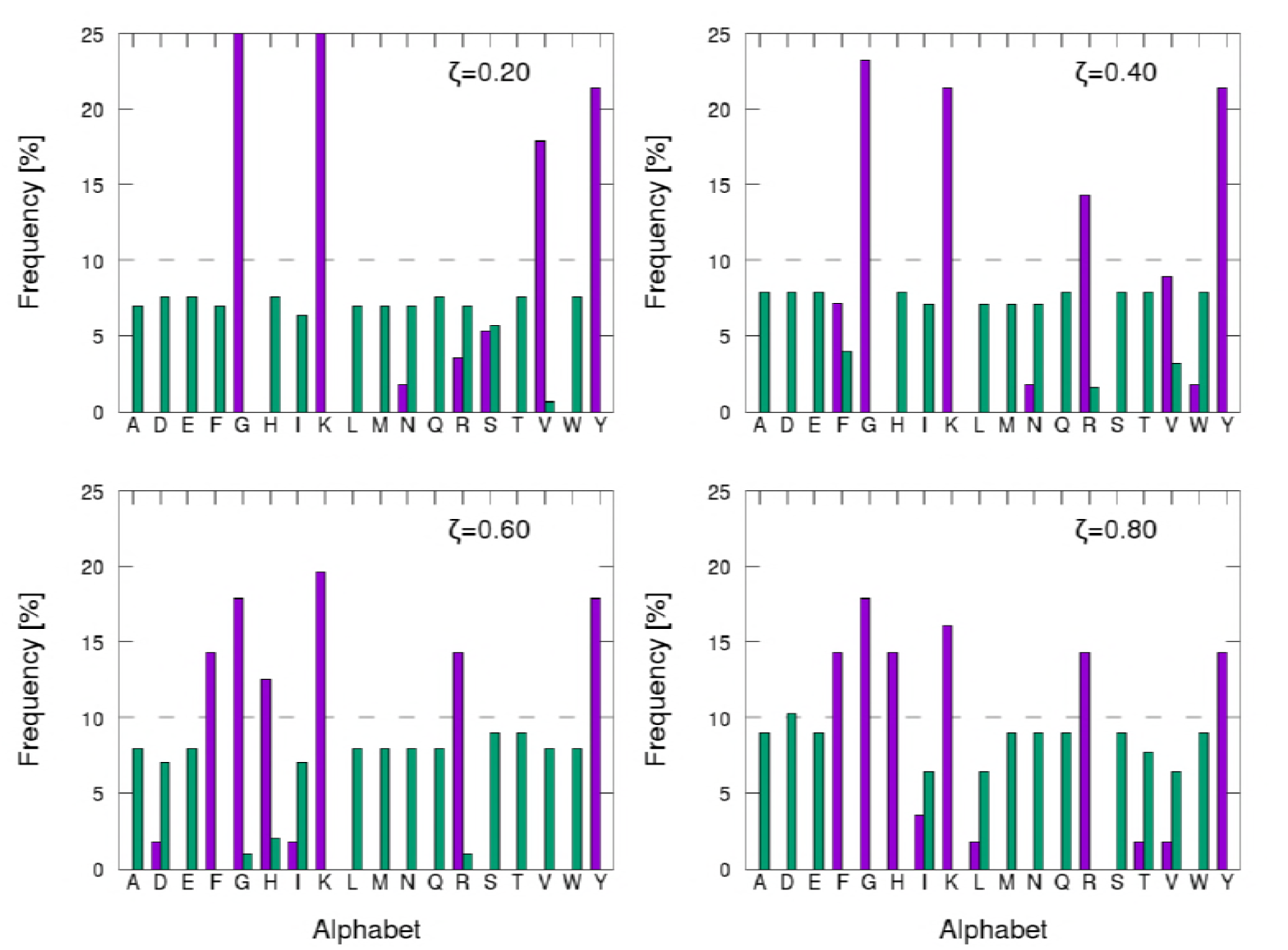
Frequency of amino acids in the designed sequences. The grey dotted line is the value used to consider one amino acid in the effective alphabet. Purple bars refer to the sequence of protein G, while green bars to the sequences of the binding site. Cysteine (C) and Proline (P) are not included in the design alphabet.

To test the selected sequences, we first examine the folding stability of the protein itself, therefore performing a folding simulation in an empty box starting from a fully stretched configuration. The set of reduced temperatures for these simulations (2.0; 1.8; 1.6; 1.4; 1.3; 1.2; 1.1; 1.0; 0.9; 0.8; 0.7; 0.65; 0.6; 0.55; 0.5; 0.45) is chosen in order to observe repeatedly folding events. Fig. 4 shows the free energy profiles as a function of *DRMSD*. From previous works (7, 9), the criterion for assessing a stable fold is to observe a funnel shape of the free energy profile and a global free energy minimum for *DRMSD* ≤ 2 Å. Using this criterion, we can say that all protein sequences fold back into the target configuration, although with different precision. Sequences with a larger effective alphabet fold with higher precision, as can be seen from the *DRMSD* value of the configurations corresponding to the global free energy minimum for each system (right hand side of Fig. 4): *DRMSD* = 2.1 Å for *ζ* = 0.20; *DRMSD* = 1.9 Å for *ζ* = 0.40; *DRMSD* = 1.3 Å for *ζ* = 0.60 and *DRMSD* = 1.5 Å *ζ* = 0.80. ^1^ The sequence optimised for the binding site *ζ* = 0.40 shows a secondary minimum in the free energy, corresponding to misfolded compact structures, therefore being the system less stable for the folding in the bulk.

**Figure 4:**
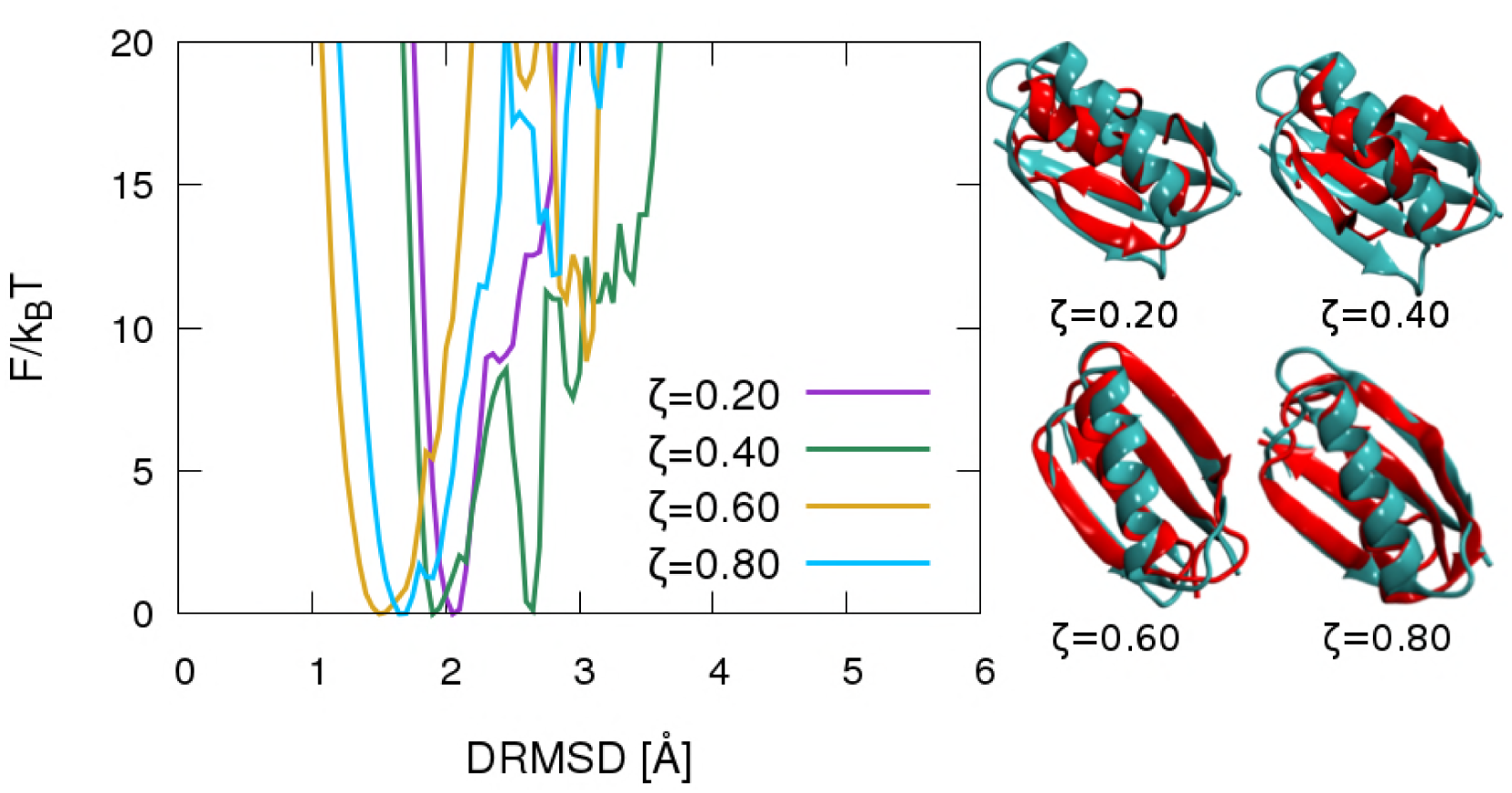
Folding free energy profiles *F/k*_*B*_*T* of single protein (no binding site) at reduced temperature 0.55 as a function of DRMSD from the native target structure (Protein G, PDB ID: 1PGB). different colours represent the folding free energy of the protein sequence obtained via the design procedure in the presence of the binding site characterised by the *ζ* value specified in the key. Configurations corresponding to the free energy minimum for each system are represented in red, compared to the native protein G (in green).

From the described scenario, we can draw two important conclusions: firstly, design with a limited alphabet of 4 letters can produce a funnel-like folding free energy landscape; secondly, with 6 letters we recover the folding precision of previous Caterpillar designs made with 20 letters (9). Our results are consistent with the experimental observation that 6 letters are a minimal set necessary to maintain protein structure and function (36, 38, 41–44).

From the Random Energy Model (49–51) we know that for a heteropolymer to be designable it has to satisfy the relation *q* > exp *ω*, where q is the alphabet size and *ω* the conformational entropy per residue. Hence, a 4 letters alphabet gives an upper bound to the conformational entropy *ω* of the Caterpillar backbone and therefore of the more restricted natural protein backbones. Such a result is compatible with the recent observations of Cardelli *et al*. (52) who mapped the designability phase space for a general heteropolymer decorated with directional interactions similar to the hydrogen bonds present along the protein backbone. For polymers with two directional interactions per particle the minimum alphabet measured was three, i.e. close to the one presented here.

To test the effect of the alphabet restriction on protein-protein interactions, we perform folding simulations in the presence of the binding site. In Fig. 5 we plot the free energy landscape as a function of *DRMSD*_*intr a*_ using the native protein G as target configuration, and *DRMSD*_*inter*_ using the folded bound configuration as a target. This choice allows us to monitor the folding and binding properties of the system independently. Conformations that are folded and bound can be found in the bottom left corner, while folded unbound ones in the top left one.

**Figure 5:**
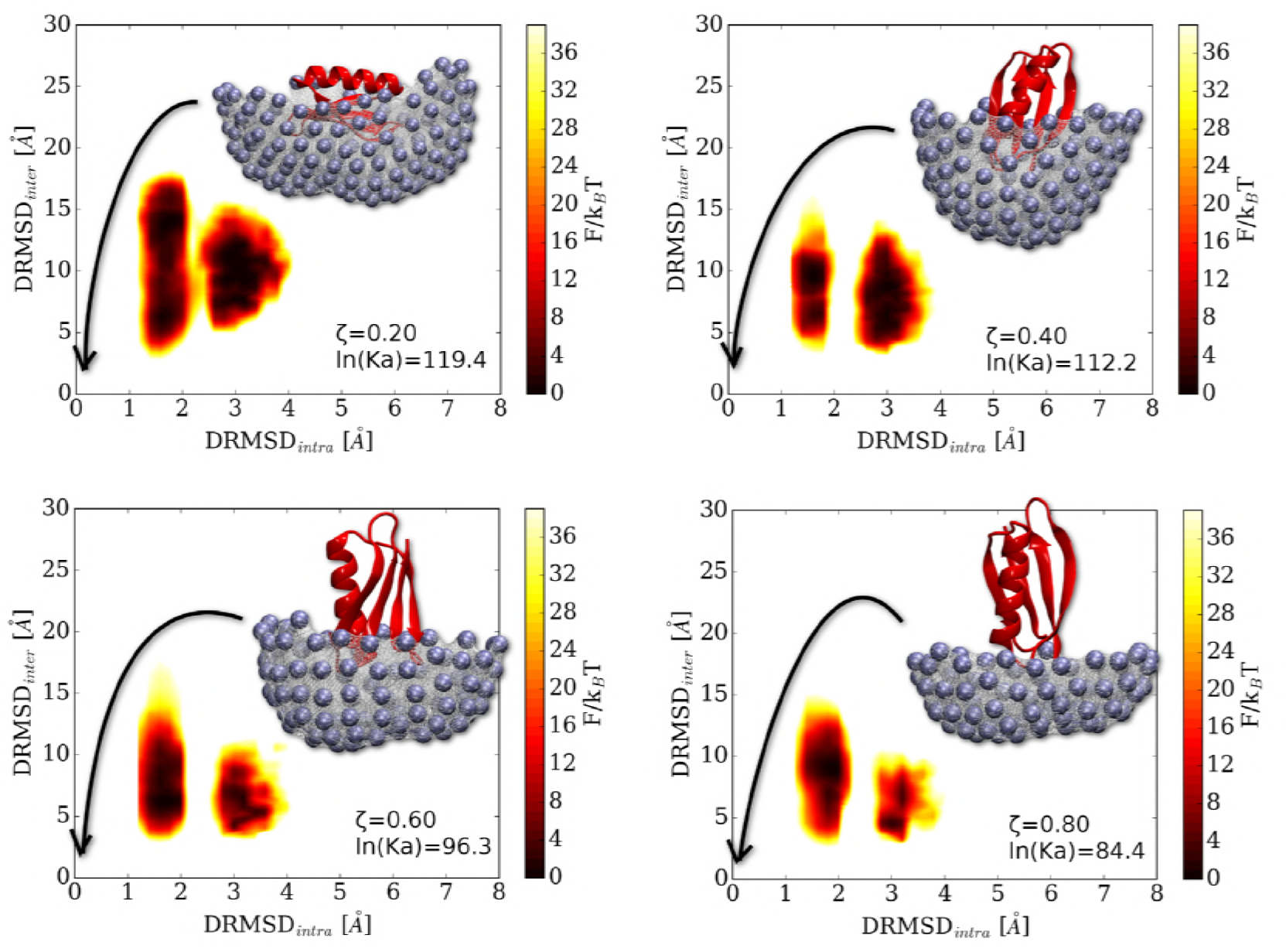
Folding free energy landscapes *F*/*k*_*B*_*T* at reduced temperature 0.76 as a function of *DRMSD*_*intr a*_ protein from the native protein G as target and *DRMSD*_*inter*_ protein-binding site from the folded protein bound to the binding site (configurations depicted in the panels). The binding affnity decreases along with the binding site size, as shown by the value of the association constants *K*_*a*_ in the plot key.

Additionally, we also check the free energy profiles as a function of *DRMSD*_*intr a*_ for bound conformations (see Fig. 6) and in bulk solution where no protein-site contacts are possible (see Fig. 7. For a sketch of the definition of interacting and bulk solution configurations see Fig. S11 in the SI. We have also verified that the free energy profiles of configurations in the latter region correctly fold into the target structure (Fig. 7), consistently to what we observe in the isolated protein folding simulations.

**Figure 6:**
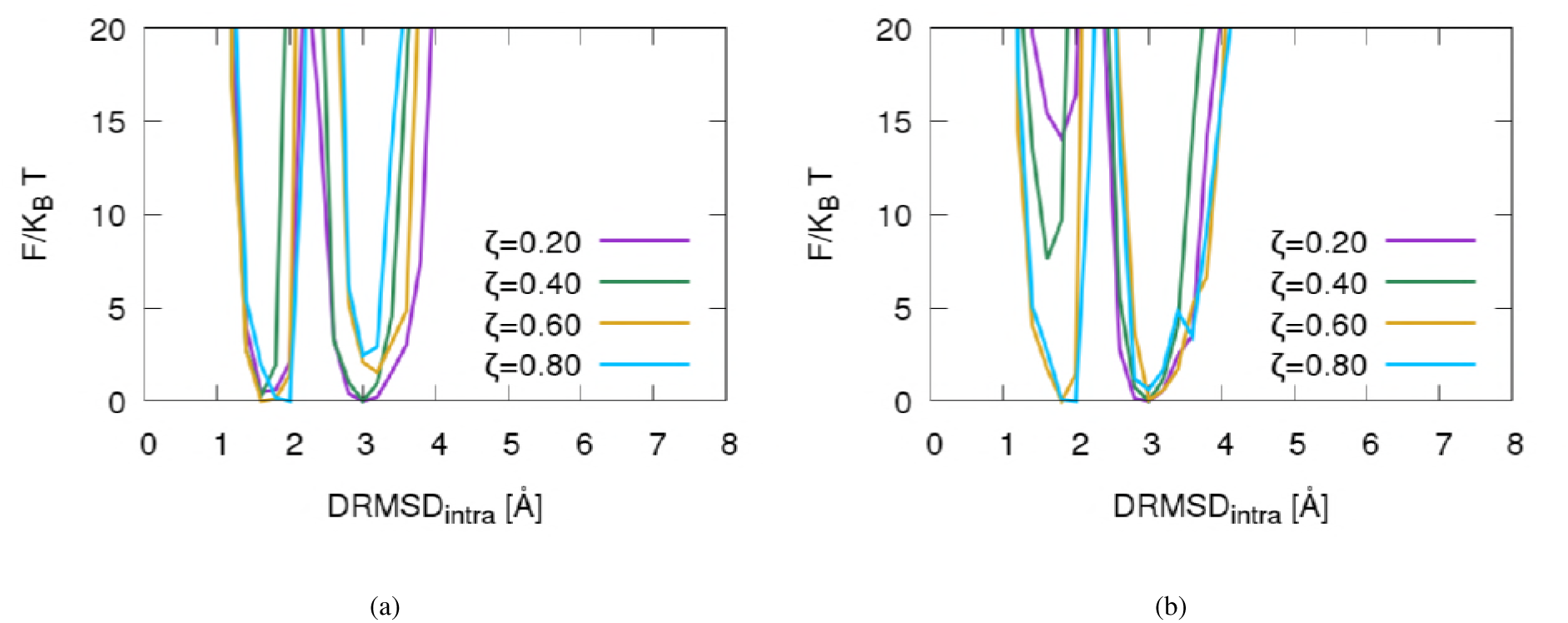
Folding free energy profiles *F*/*k*_*B*_*T* at reduced temperature 0.76 (left panel) and 1.00 (right panel) as a function of DRMSD intra protein from the native target structure. Simulations of protein G in presence of binding site at *ζ* = (0.20, 0.40, 0.60, 0.80). The curves have been evaluated on bound configurations, i.e. where the protein is directly interacting with the binding site. Left panel: The presence of two equivalent minima suggests that the protein binds in the target configuration as well as in a misfolded state with the same probability. Right panel: upon increasing the temperature, the correctly folded state is destabilised for large binding sites.

**Figure 7:**
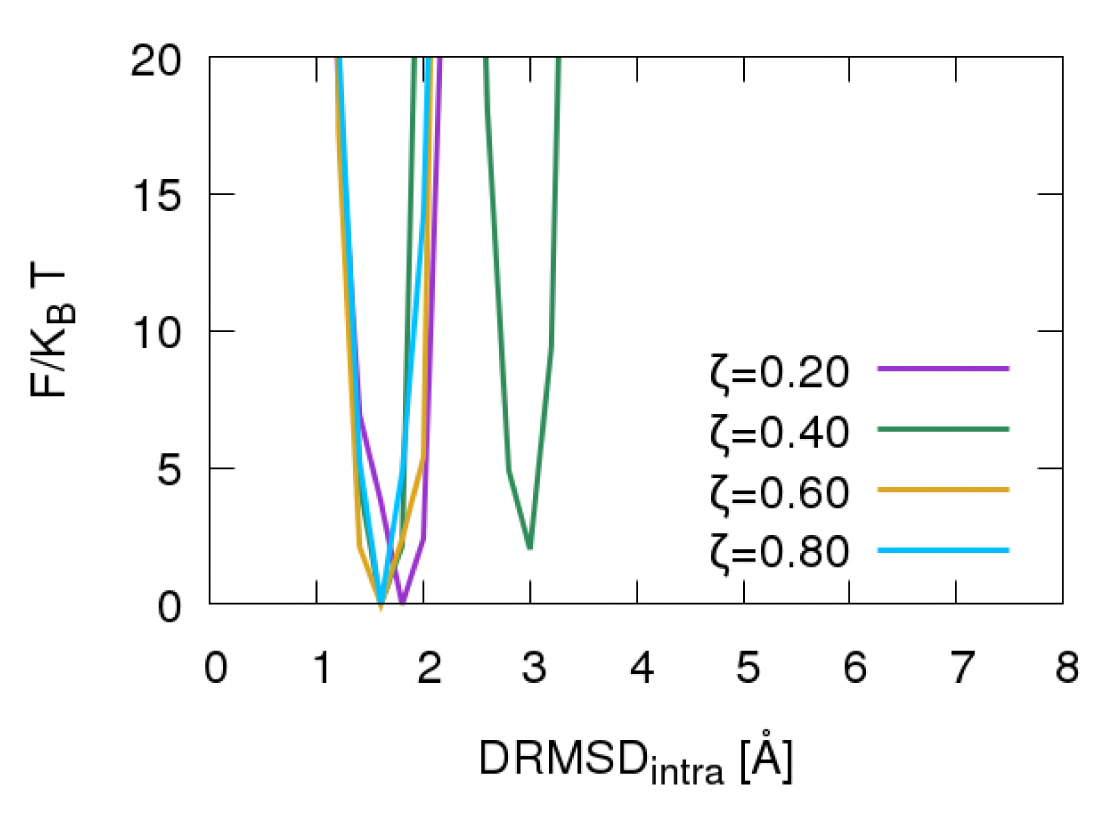
Folding free energy profiles *F* /*k*_*B*_*T* at reduced temperature 0.76 as a function of *DRMSD*_*intr a*_ protein from the native target structure. Simulations of protein G in presence of binding site at *ζ* = (0.20, 0.40, 0.60, 0.80). The curves have been evaluated on configurations in the bulk solution, where the protein-binding site distance is larger than the interaction one. The presence of a minimum below *DRMSD* =2 Å shows that the protein is folded when bound.

For all the scenarios, upon binding to the site, we observe a significant enhancement of the stability of the misfolded configurations with respect to the bulk solution (compare Fig.s 5, 6(a) and 7). In particular, there is a considerable shift in the equilibrium towards states at *DRMSD* ∼ 3 Å that are now comparable in free energy to the target configurations. This effect is due to the small protein alphabet that cannot now encode heterogeneous patterns to reduce the binding interaction with the large binding sites. It should be noted that natural binding sites expose much smaller surface areas then the one modelled here. Hence, it might be that the effect is mitigated when real protein binding sited are considered.

Analysing the behaviour of the binding process as a function of temperature we find that the random binding is so strong that the van’t Hoff curve (53, 54) shows an exothermic process above the folding temperature with positive binding affinity (Supplementary Fig. S12; for details about the evaluation of the association constant see Fig. S11). At the same time the equilibrium shifts from partially-misfolded to fully-misfolded, indicating that the unfolding process takes place at the surface while the protein remains bound (see Fig. 6(b)). Considering the large surface area and the small alphabet employed it is not surprising that specificity of the protein-protein interactions could not be achieved. This is an essential factor that could explain the size of natural alphabets, i.e. the 20 amino acids. The study of the effect of the size of the binding site on the specificity is the object of upcoming work.

## CONCLUSION

The design procedure employed in our work has a significant segregation effect on the alphabet letters used in the protein sequence. The larger the number of residues on the binding site, the smaller is the effective alphabet available for the protein sequence. On the one side, the design is capable of selecting a subset of letters that still allows the folding of the protein in the bulk solution even for the smallest effective alphabet (4 letters). The precision of the folding increases with the effective alphabet size. Interestingly, the experimentally determined alphabet size of 6 letters is also what we observed recovering the design accuracy commonly obtained with a 20 letter alphabet. Finally, the identified alphabets in this work are automatically optimised by the design and are composed of the most attractive residues while maintaining the broadest heterogeneity of the solvent interactions.

Our results have far-reaching implications both in the field of protein design and for the understanding of protein evolution. In protein design, the possibility of using a reduced alphabet would considerably accelerate the search of the sequence space for good folders. In the field of protein evolution instead, the understanding of the smallest alphabet necessary for accurate proteins design is still an open question. To the best of our knowledge, this study represents the first successful design of a full natural protein structure with a reduced alphabet of just 4 letters.

## AUTHOR CONTRIBUTIONS

I.C. and C.D. designed the research; F.N. and L.T. developed the simulation tools; F.N. performed the simulations and the data analysis. All the authors discussed the research and participated in the writing and the revision of the article.

## ACKNOWLEDGEMENTS

All simulations presented in this paper were carried out on the Vienna Scientific Cluster (VSC). We acknowledge support from the VSC School, as well as from the Austrian Science Fund (FWF) project 26253-N27. V. B. acknowledges the support from FWF Grant No. M 2150-N36.

## SUPPLEMENTARY INFORMATIONS

### Binding site modelling

The artificial binding site is a mould obtained by pushing the folded protein conformation on a planar mesh of self avoiding beads (self avoiding radius = 2 Å).

To obtain the mould, we initially generate a dense mesh in the *z* = 0 plane by placing the beads on a 2D square lattice with step 0.5 Å.

We then identify the centre of mass of the protein and use it to measure the height of the protein with respect to the plane, which is one of our main control parameters. Specifically, we consider the CM height *z*_*C M*_ normalised over the maximum distance,*r*_*M AX*_, of a *C*_*α*_ from the CM of the protein, so that 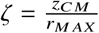 ranges between 0 (when the CM lies on the *z* = 0 plane) and 1.

We consider several values of *ζ* and for each of them we identify a protein orientation which maximise the contact surface with the binding site according to the following procedure. We begin by aligning the minor inertia axis of the protein with the *z* axis and then perform a discrete set of rotations around the *x* and *y* axes. For each of these rotations, we recompute the mould, map it to a triangular mesh, and calculate the surface of the latter. The area of the surface is obtained by summing the area of each triangle formed by all the triplets of the mesh points at *z ≠* 0.

The binding site is obtained by setting a minimum protein-mesh bead distance *µ* = 13 Å and using it to inflate the radius of the protein *C*_*α*_s, thus generating a mould by pushing downwards all mesh beads within the inflated radius. To avoid large gaps within the binding site, we perform an iterative smoothing procedure. For each bead of the pocket we compute its distance to all its neighbouring beads on the square lattice, and if this is larger than the self-avoiding radius, we shift the vertical position of the bead so to fill the gap.

In order to confer a protein nature to the artificial binding site, a homogeneously distributed set of mesh points are “activated”, i.e. switched from pure self avoiding beads to *C*α atoms. In order to do so, we firstly switch all the beads of the mesh to *C*_*α*_, then we set a minimum binding site *C*_*α*_ – *C*_*α*_ distance *δ* = 5 Å and loop over all bead points starting from a corner of the mesh. For each activated atom in the loop we deactivate all points within a sphere of radius *δ*. Clearly, after the first atom the loop will incur both in activated and de-activated beads. The latter are simply skipped by the procedure. Finally, we deactivate all the beads at *z* = 0, so that only the pocket contains *C*_*α*_ beads.

---

**Figure S8:**
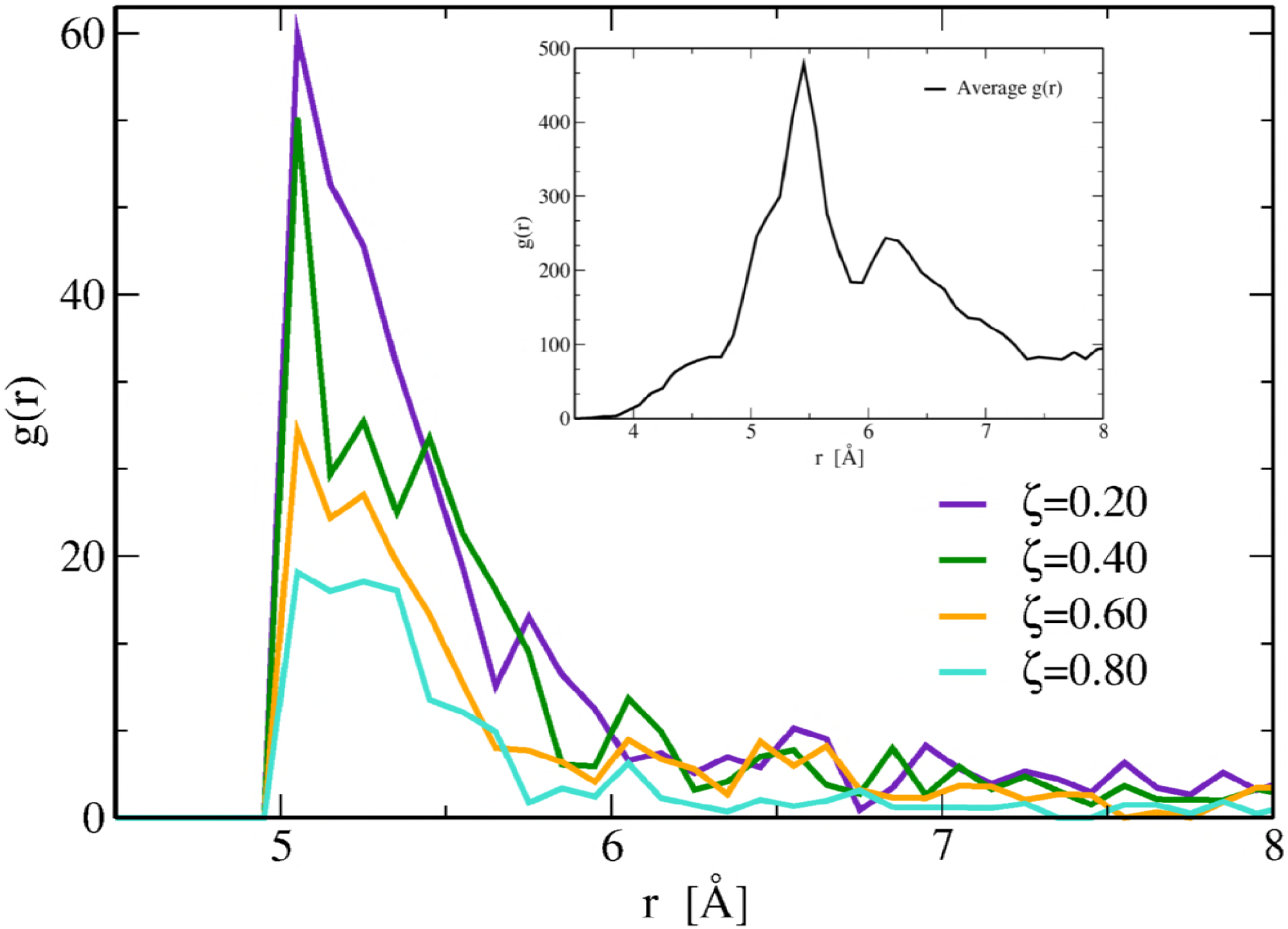
Radial distribution function *g r* calculated over the active residues *C*_Surf_ of the binding site placed at the minimum distance *δ* = 5 Å from each other. The latter was determined by identifying the value of *δ* that yields the mean nearest distance 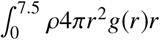 closest to 5.7 Å obtained by averaging over 145 protein structures taken from the PDB (black line in the inset). The integral interval was chosen visually taking a region large enough to fully contain the first peak in the *g*(*r*). It is important to stress that in the protein *g r* the selected distribution includes the contribution from all secondary structure elements (52).

**Figure S9:**
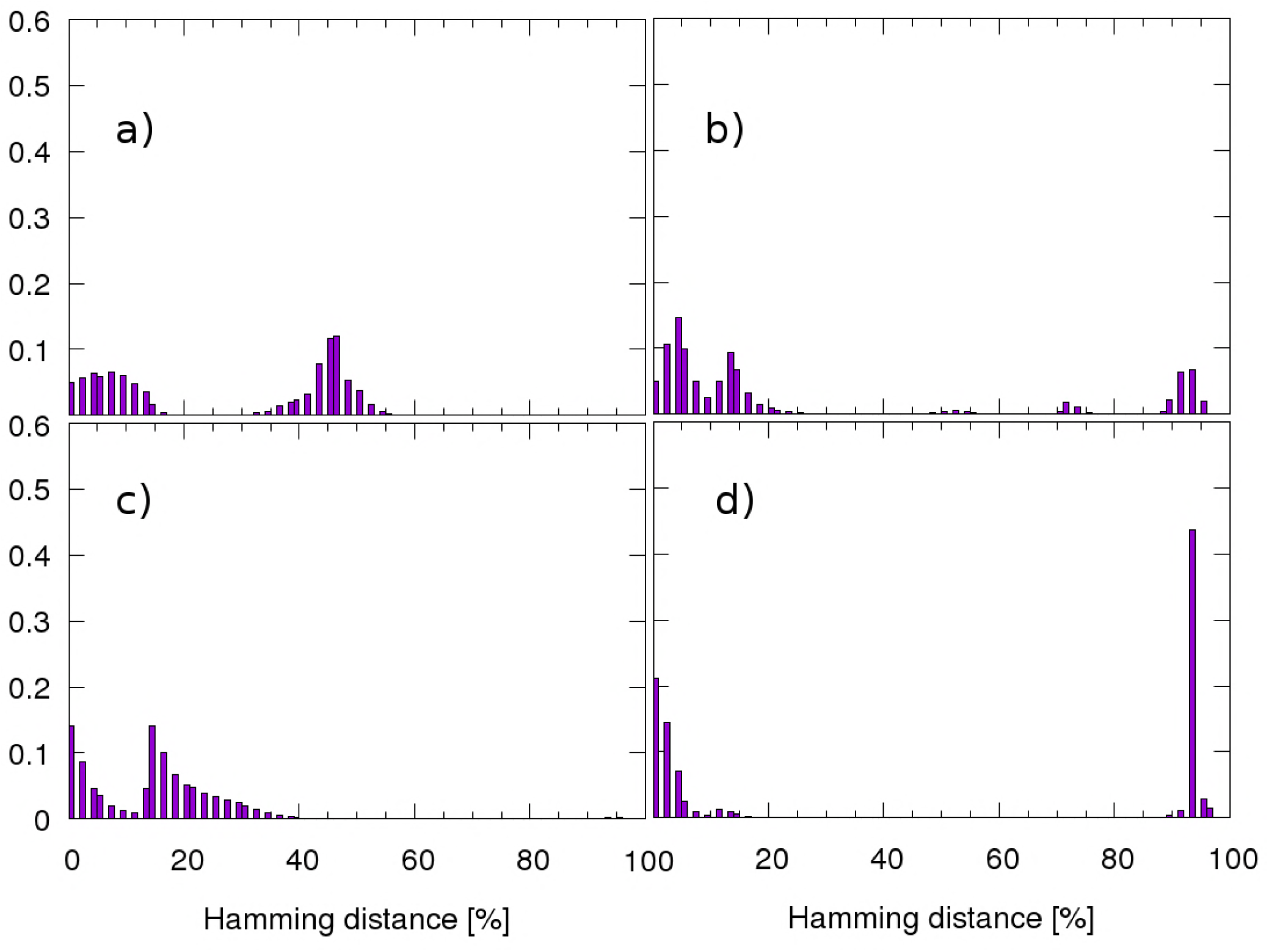
Histograms obtained by evaluating the Hamming distance (% relative to the chain length) between all possible pairs of sequences chosen selecting 200000 solutions around the design free energy minimum of systems *A* and *B*, corresponding to the *ζ* values specified in the following. All panels refer to self comparison of sequences belonging to the same basin. a) *A*: *ζ*= 0.20; *B*: *ζ*= 0.20 b) *A*: *ζ*= 0.40; *B*: *ζ*= 0.40 c) *A*: *ζ*= 0.60; *B*: *ζ*= 0.60 d) *A*: *ζ*= 0.80; *B*: *ζ*= 0.80

**Figure S10:**
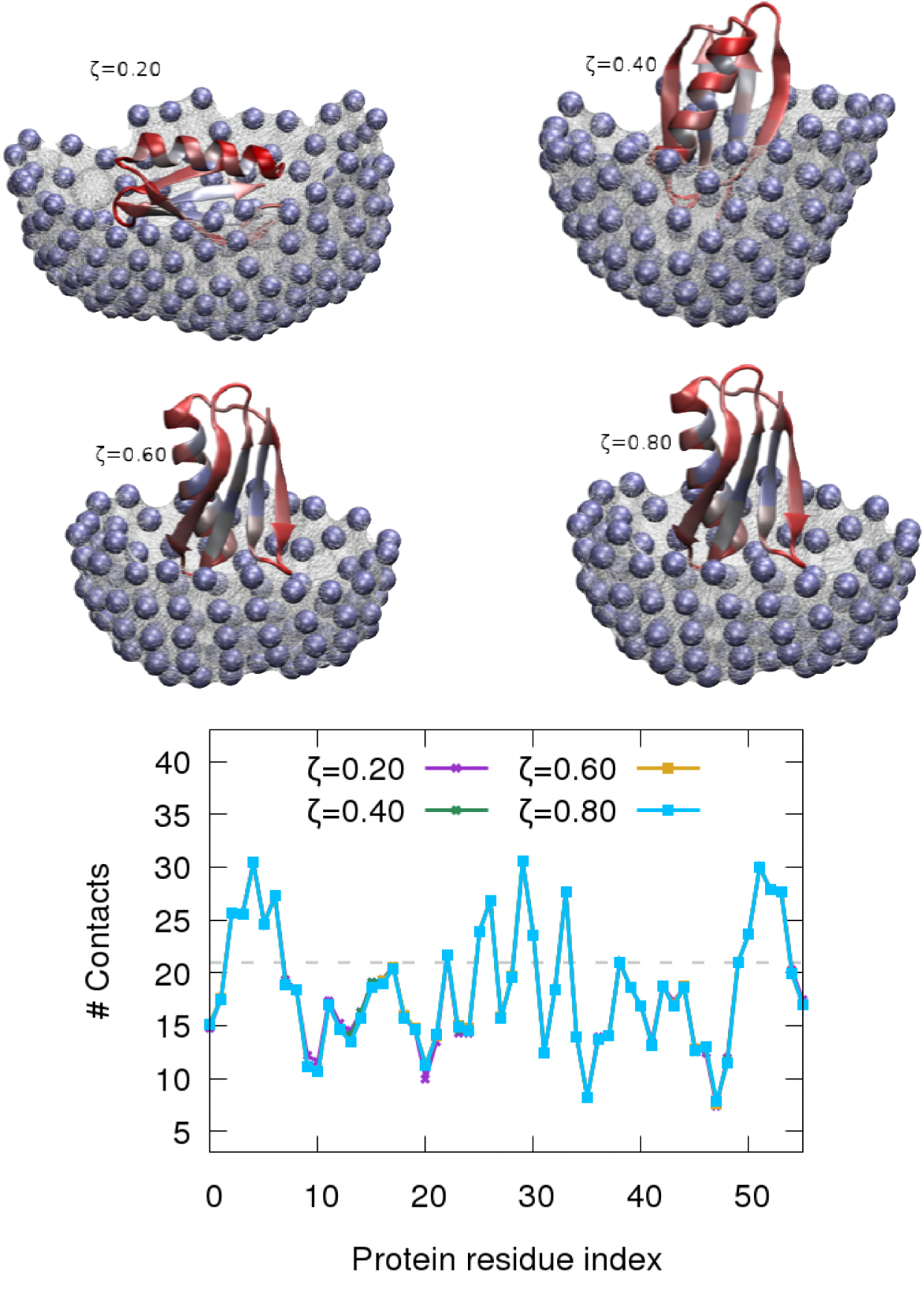
Water exposure profiles of protein G in presence of binding sites constructed with different *ζ* values. The colour scheme of the protein is related to the number of total contacts: solvent exposed regions in red, buried in blue. The plot shows the number of contacts for each protein residue. The grey dashed line defines the threshold Ω = 21 on the number of neighbours above which an amino acid is considered buried in the protein core. We used the information relative to the exposure of protein G to set the radius of the probe sphere used in the evaluation of the number of contacts for each active *C*_*α*_ of the binding site (the blue spheres in the figure). We counted the offset for the number of neighbours for each of the binding site *C*_*α*_ by multiplying the fraction of the volume of a sphere of radius 6 Å that lies under the mesh surface by the average amino acid density in globular proteins (ρ = 0.011687 *aa /* Å^3^, evaluated over a set of 145 globular proteins). The radius of the sphere was optimized to reach an average exposure of each residue similar to the one of the most exposed residues of protein G (corresponding to the minima in the plot).

**Figure S11:**
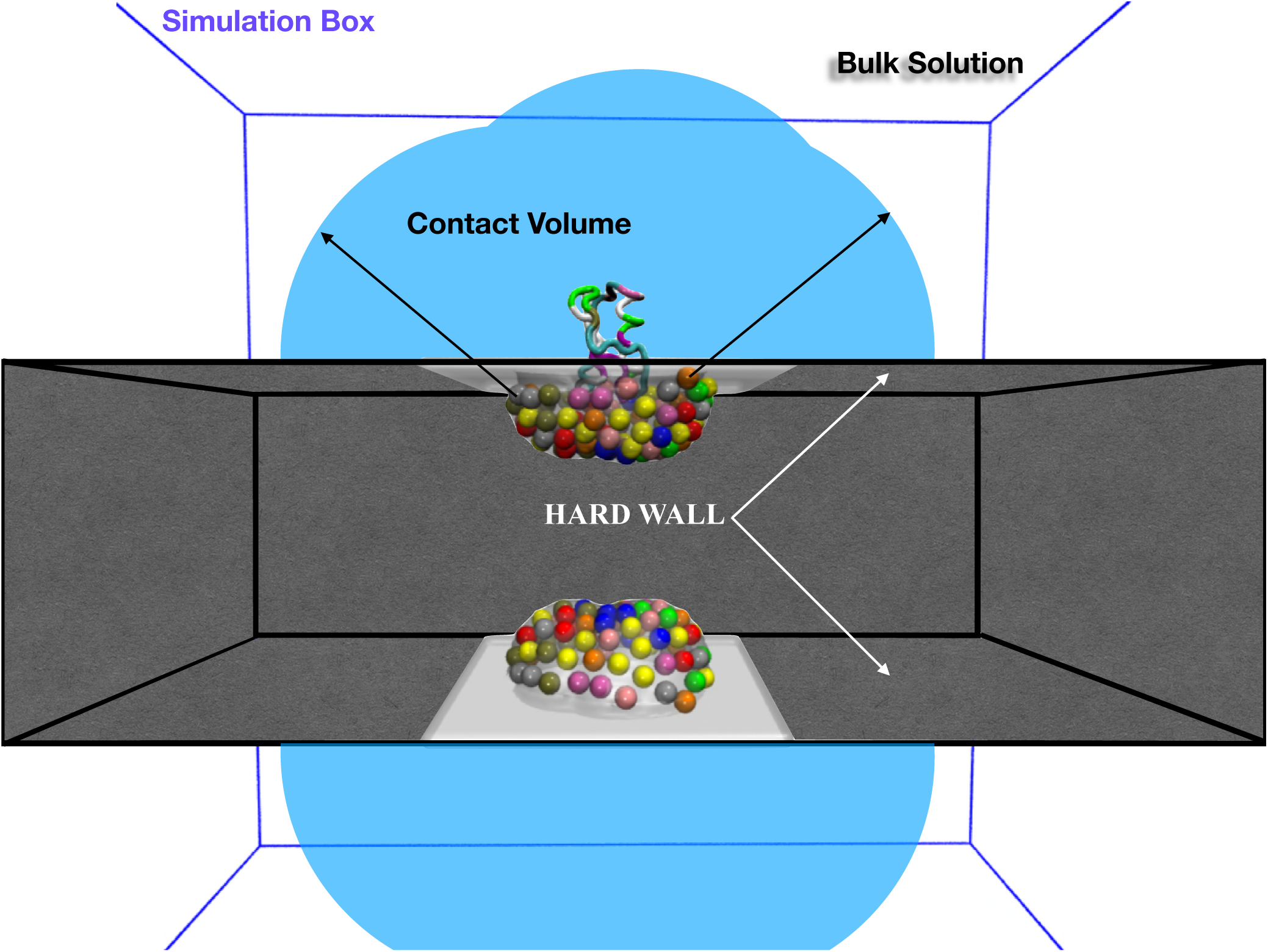
Simulation box for protein folding in presence of two copies of a potential partner. The partner is depicted as a binding site surface decorated with *C*α atoms, i.e. the coloured spheres in the image. Each colour, both on the protein backbone and on the *C*α of the binding sites, represents a different amino acid type. The protein sequence and the identity of the binding site amino acids are designed simultaneously with the procedure described in the `Design` sub-section. The protein is a flexible polymer diffusing in the box under Periodic Boundary Conditions. A mirroring move provides a swap between opposite chiralities, since the Caterpillar model does not take into account the amino acid chirality. The cubic box contains two binding sites, one the mirror image of the other, the position of which is fixed during the simulation. The binding sites are far apart enough to prevent the protein to interact with both at the same time. A hard wall between the binding sites prevents the protein to approach the surfaces from the convex side. For the evaluation of the association constant 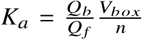, we need to assess the number of binding sites in the box *n* = 2, define the accessible simulation volume *V*_*box*_ and discern bulk solution configurations (contributing to *Q* _*f*_, the partition function of the free protein) from the ones where the protein is directly in contact with the binding site (contributing to *Q*_*b*_, the partition function of the bound configurations). The accessible volume is easily obtained by subtracting the hard wall contribution to the box volume. The bound configurations are the ones characterised by a protein-binding site interaction energy *E*_*inter*_ ≠ 0. The configurations contributing to *Q* _*f*_, instead, are the ones for which the central amino acid of the protein is at a distance *r* > *H*_*lenght*_ + *R*_*int*_ with respect to all the amino acids of the surface (being *H*_*lenght*_ half of the stretched protein length; *R*_*int*_ the interaction radius) and, therefore, where the interaction with the binding site is not possible. We refer to the latter region as the bulk solution.

**Figure S12:**
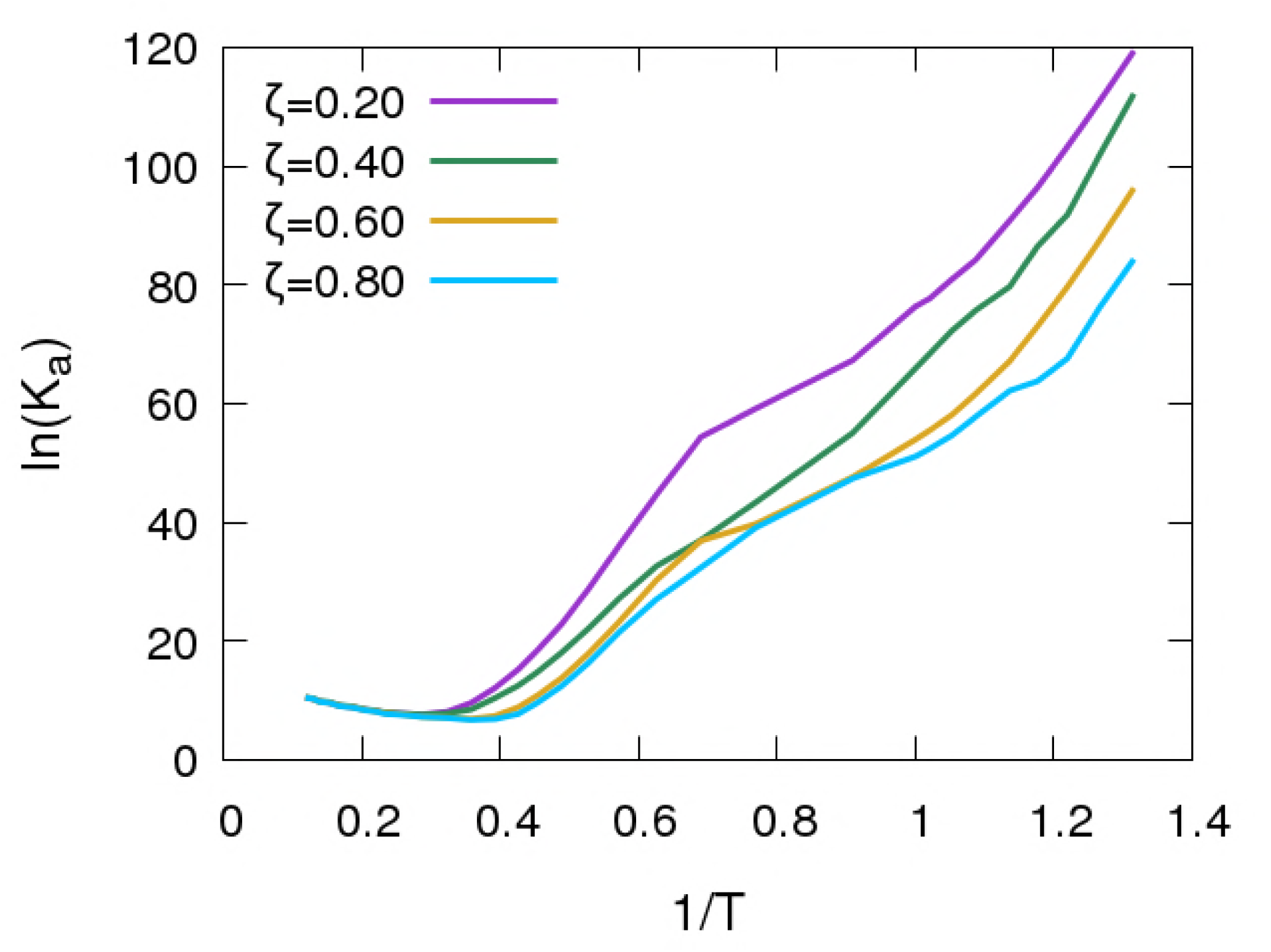
Van’t Hoff plot of ln *K*_*a*_ as a function of the inverse reduced temperature 1 *T* for the investigated systems. The association constant *l mol* is computed as *K*_*a*_ = exp Δ*F k*_*B*_*T V*_*box*_ *n*, where *V*_*box*_ is the accessible volume of our simulation box, *n* = 2 is the number of binding sites and Δ*F* = − *k*_*B*_*T* ln *Q*_*b*_ *Q* _*f*_ is the binding free energy (29). The partition functions *Q*_*b*_ and *Q* _*f*_ refer to all protein conformations bound to the binding site and free in the bulk solution, respectively.

## Footnotes

1 The DRMSD values correspond to 4.9; 5.5; 2.4 and 2.7 Å in *RMSD* respectively.

